# Real-time imaging of *Arc/Arg3.1* transcription *ex vivo* reveals input-specific immediate early gene dynamics

**DOI:** 10.1101/2021.12.16.472958

**Authors:** Pablo J. Lituma, Robert H. Singer, Sulagna Das, Pablo E. Castillo

## Abstract

The ability of neurons to process and store salient environmental features underlies information processing in the brain. Long-term information storage requires synaptic plasticity and regulation of gene expression. While distinct patterns of activity have been linked to synaptic plasticity, their impact on immediate early gene (IEG) expression remains poorly understood. The activity regulated cytoskeleton associated (*Arc*) gene has received wide attention as an IEG implicated in synaptic plasticity and memory. Yet, to date, the transcriptional dynamics of *Arc* in response to compartment and input-specific activity is unclear. By developing a knock-in mouse to fluorescently tag *Arc* alleles, we studied real-time transcription dynamics after stimulation of dentate granule cells (GCs) in acute hippocampal slices. To our surprise, we found that *Arc* transcription displayed distinct temporal kinetics depending on the activation of excitatory inputs that convey functionally distinct information, i.e. medial and lateral perforant paths (MPP and LPP, respectively). Moreover, the transcriptional dynamics of *Arc* after synaptic stimulation was similar to direct activation of GCs, although the contribution of ionotropic glutamate receptors, L-type voltage gated calcium channel, and the endoplasmic reticulum (ER) differed. Specifically, we observed an ER-mediated synapse-to-nucleus signal that supported elevations in nuclear calcium, and rapid induction of *Arc* transcription following MPP stimulation. However, activation of LPP inputs displayed lower nuclear calcium rise, which could underlie the delayed transcriptional onset of *Arc*. Our findings highlight how input-specific activity distinctly impacts transcriptional dynamics of an IEG linked to learning and memory.

**Significance statement:** Environmental experiences trigger neuronal activity that elicits gene expression in the nervous system. Rapid induction of specific genes known as immediate early genes (IEGs) supports activity-dependent changes of neuronal circuits to ultimately influence animal behavior. However, the cellular and molecular mechanisms controlling how distinct forms of neuronal activity modulate IEG expression remains unclear. The activity regulated cytoskeleton associated (*Arc*) gene is a critical IEG linked to memory. By imaging *Arc* transcription in real-time after neuronal activity, we identified how different receptors and signaling pathways influence transcriptional induction and dynamics of an IEG. Our findings provide insights into how information received by distinct synaptic inputs could be encoded by modulating IEG dynamics.

## Introduction

Encoding of salient stimuli through changes in immediate early gene (IEG) expression highlights a crucial aspect of information processing in the brain (1–3). The activity regulated cytoskeleton associated (*Arc*) gene has received wide attention as a critical IEG involved in certain forms of memory (4–6). Transcriptional activation of *Arc* occurs after long-term changes in neurotransmission (7–9) and the *Arc* mRNAs localize to recently activated synapses (7). Local synthesis of Arc protein was suggested to modulate synaptic strength by stabilization of F-actin and endocytosis of AMPA receptors (AMPARs) (10, 11). Therefore, Arc impacts different forms of synaptic plasticity: long-term potentiation and long-term depression (5, 11–14), but see (15), as well as homeostatic synaptic scaling (16). In addition to influencing synaptic functions, evidence for Arc protein regulation of plasticity related genes is emerging (17, 18). Taken together, these studies emphasize that precise tuning of Arc protein levels is required to support synaptic plasticity and long-term memory. Thus, elucidating the mechanisms controlling the temporal dynamics of *Arc* gene expression is crucial to our understanding of information processing in the brain.

The pattern and location of neuronal activity required for IEG expression and temporal dynamics remains an open question (19–21). Most studies evaluating how neuronal activity is transduced into gene transcription are primarily based on genome-scale analyses (22) or mRNA detection in fixed tissue (7, 19, 23). Modified *in situ* hybridization (ISH) approaches identified *Arc* transcribing neurons after a behavioral task or synaptic stimulation (7, 23). In recent years, *Arc* transcribing neurons have been tracked using reporters expressing GFP or dVenus driven by IEG promoters (24–26). However, visualizing the complexity of endogenous *Arc* transcriptional dynamics remains challenging in live tissue. Overcoming such limitation is important for studying how neurons distinctly encode environmental features through changes in IEG expression. To address this unknown and advance gene-imaging technology, we developed a conditional Cre recombinase approach that fluorescently labels endogenous *Arc* mRNAs in mice. Our tagging strategy was able to detect allelic *Arc* transcription in tissue with high temporal sensitivity. Since transcription is the first step of IEG regulation, imaging Arc transcription kinetics is a good indicator of how activity is transduced into different temporal patterns of Arc expression.

Using acute hippocampal slices from our reporter mouse, we measured the magnitude and temporal kinetics of *Arc* transcription after direct activation of dentate granule cells (GCs) and selective activation of the excitatory synaptic inputs medial perforant path (MPP) and lateral perforant path (LPP). We found that presynaptic patterns of activity led to distinct temporal dynamics of *Arc* transcription. MPP synapses rapidly and robustly activated *Arc* transcription via a synapse-to-nucleus signal supported by the ER that contributes to nuclear calcium (Ca^2+^) elevation. In contrast, LPP stimulation resulted in delayed *Arc* transcription and lower nuclear Ca^2+^ rise. Furthermore, we developed an optical stimulation method to trigger *Arc* transcription in GCs and identify the relative contributions of receptors and subcellular signaling in optical versus synaptic driven neuronal activity. Our findings strongly suggest that excitation-transcription coupling of IEGs in neurons display input-specificity, and that compartmentalized neuronal activity may encode specific environmental features in the genome.

## Results

### Visualizing real-time transcription of endogenous *Arc* in live tissue after neuronal stimulation

To monitor gene transcription in real time, we utilized the stem loop technology, whereby mRNAs tagged with multimerized RNA stem loops are detected after synthesis by high-affinity binding of fluorescently labeled coat proteins (27). The endogenous *Arc* gene was tagged by introducing 24 repeats of PBS loops into the 3’UTR (28) that are bound by the coat protein PCP fused to GFP. A new mouse line where PCP-GFP expression was designed to be Cre-dependent using loxP sites and enabled cell-specific PCP-GFP labeling (Fig. 1A). This PCP-GFP mouse was then crossed with the PBS-tagged Arc mouse (Arc^P/P^), to generate double homozygous mice (Arc^P/P^ x PCP-GFP) for imaging *Arc* mRNA synthesis in live tissue (Fig. 1A). Viral delivery of Cre to hippocampal GCs elicited recombination for fluorescent labeling of *Arc* mRNAs. To induce *Arc* gene expression, stimulation of MPP and LPP synapses was confirmed by paired-pulse properties of field excitatory postsynaptic potentials (Fig. 1B). Meanwhile, optogenetic activation of the fast kinetics channelrhodopsin variant ChIEF (29) expressed in GCs elicited action potentials (Fig. 1B). Using two-photon microscopy, *Arc* transcription was detected as bright fluorescent foci in GC nuclei after neuronal stimulation (Fig. 1C, Movie S1). The emergence of fluorescent foci (intensity >10% from nuclear background) indicated *de novo* transcription of an *Arc* allele (Fig. 1C). Meanwhile, basal transcription can be maintained/enhanced by increasing the amplitude of the pre-existing allele, or by the appearance of the second allele (Fig. 1C). Quantification of these transcriptional signatures was designated as total percentage of transcribing cells in the imaging field (Fig. 1C). As a result, we could interrogate how GC activity induces *Arc* transcription *ex vivo* in real-time with high spatiotemporal resolution.

**Fig. 1.**
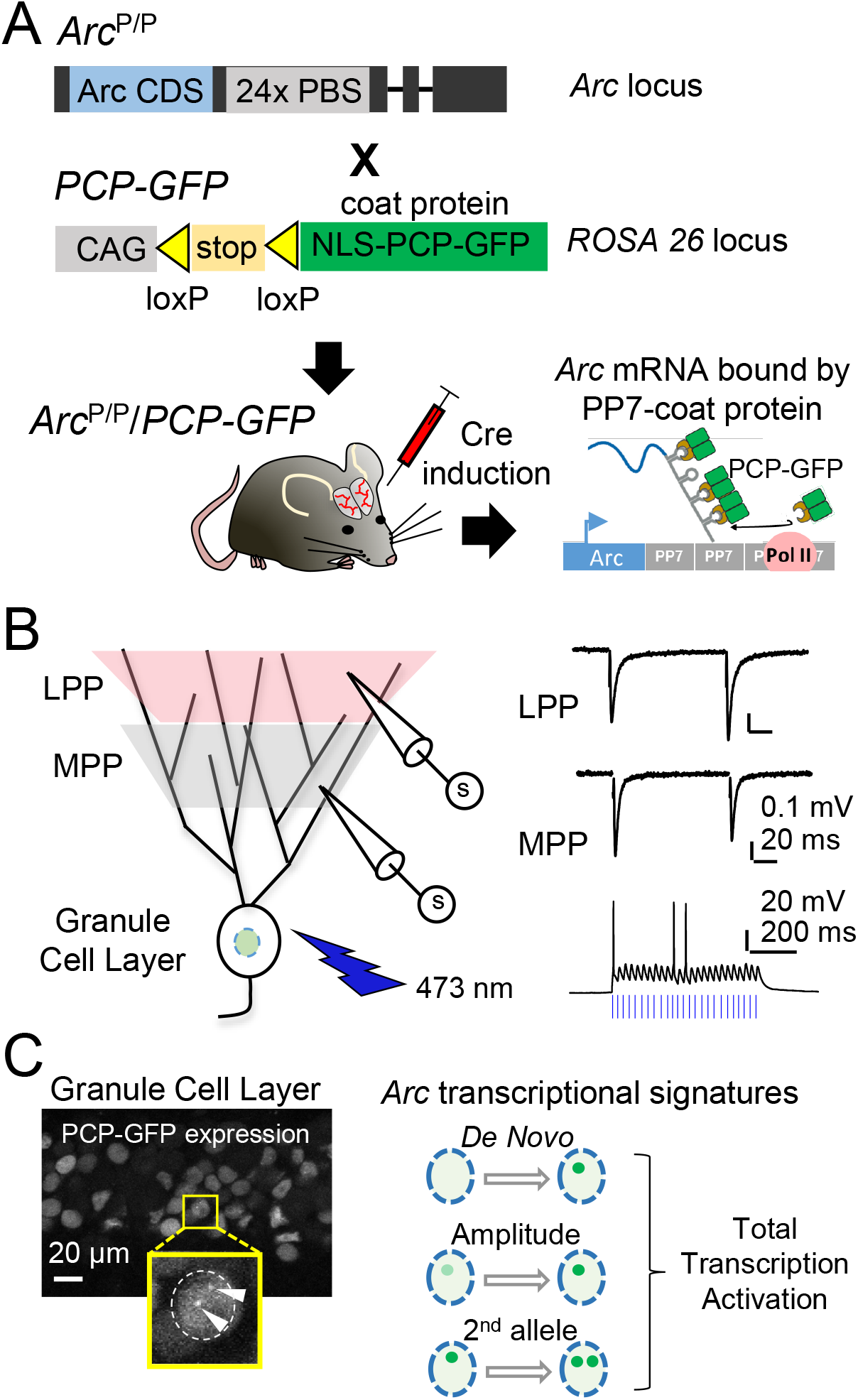
Cre-inducible *Arc* mRNA PCP-GFP tagging system to visualize transcription triggered by neuronal activity in live tissue. (***A***) *Arc*^P/P^ animals were crossed with *PCP-GFP* transgenic animals that contained PCP-GFP in the *ROSA 26* locus flanked by a stop/lox site rendering PCP-GFP expression Cre-dependent. The double homozygous mouse (Arc^P/P^/ PCP-GFP) were stereotactically injected with Cre, enabling PCP-GFP binding to the PP7 stem loops on the *Arc* mRNAs, and detection of the transcribing alleles. (***B***) Experimental arrangement for the electrical stimulation of LPP and MPP synaptic inputs and optical activation of granule cells expressing ChIEF (*left).* MPP and LPP field excitatory postsynaptic potentials elicited by paired-pulse stimulation (*top, right*), and light-induced action potentials in GCs (*bottom, right*). (***C***) Representative image shows the two transcribing alleles producing tagged *Arc* mRNAs in GCs *(left).* Transcription signals were classified as emergence of new (*De Novo*) or increase of basal transcription by increase in signal intensity (Amplitude) or induction from the second allele. The combined quantification of all signals is denoted as total transcriptional activation.

### Stimulation of proximal and distal GC synapses activates *Arc* with different temporal kinetics

Long-term changes in neurotransmission and epileptic forms of activity at perforant path inputs elicit *Arc* transcription in GCs (7–9). However, the precise impact of perforant path synapses on the temporal kinetics of *Arc* transcription in GC nuclei is unknown. This is a relevant question because these inputs convey functionally distinct information; while proximal MPP inputs encode contextual information, distal LPP inputs carry the content of an experience (30–33). Hence, we determined how stimulation of MPP or LPP inputs regulates *Arc* transcriptional dynamics in real time (Fig. 2A). MPP activation significantly induced *Arc* in 12.5 ± 0.6 % of the GC population at 15 min and peaked at 30 min (Fig. 2B). Conversely, distal LPP stimulation led to *Arc* upregulation only at later time points (Fig. 2B: LPP 8.3 ± 0.9 % at 45 min; LPP 9.2 ± 1.1 % at 60 min). Unstimulated naïve slices did not exhibit any increase in *Arc* transcription (Fig. 2B: Basal naïve 2.6 ± 0.5 % at 30 min). To confirm that fluorescent foci reported mRNA synthesis, we imaged slices in the presence of the transcriptional inhibitor DRB (100 μM). Under this condition, fluorescent foci were absent and a decline in total transcribing cells was observed (Fig. S1).

**Fig. 2.**
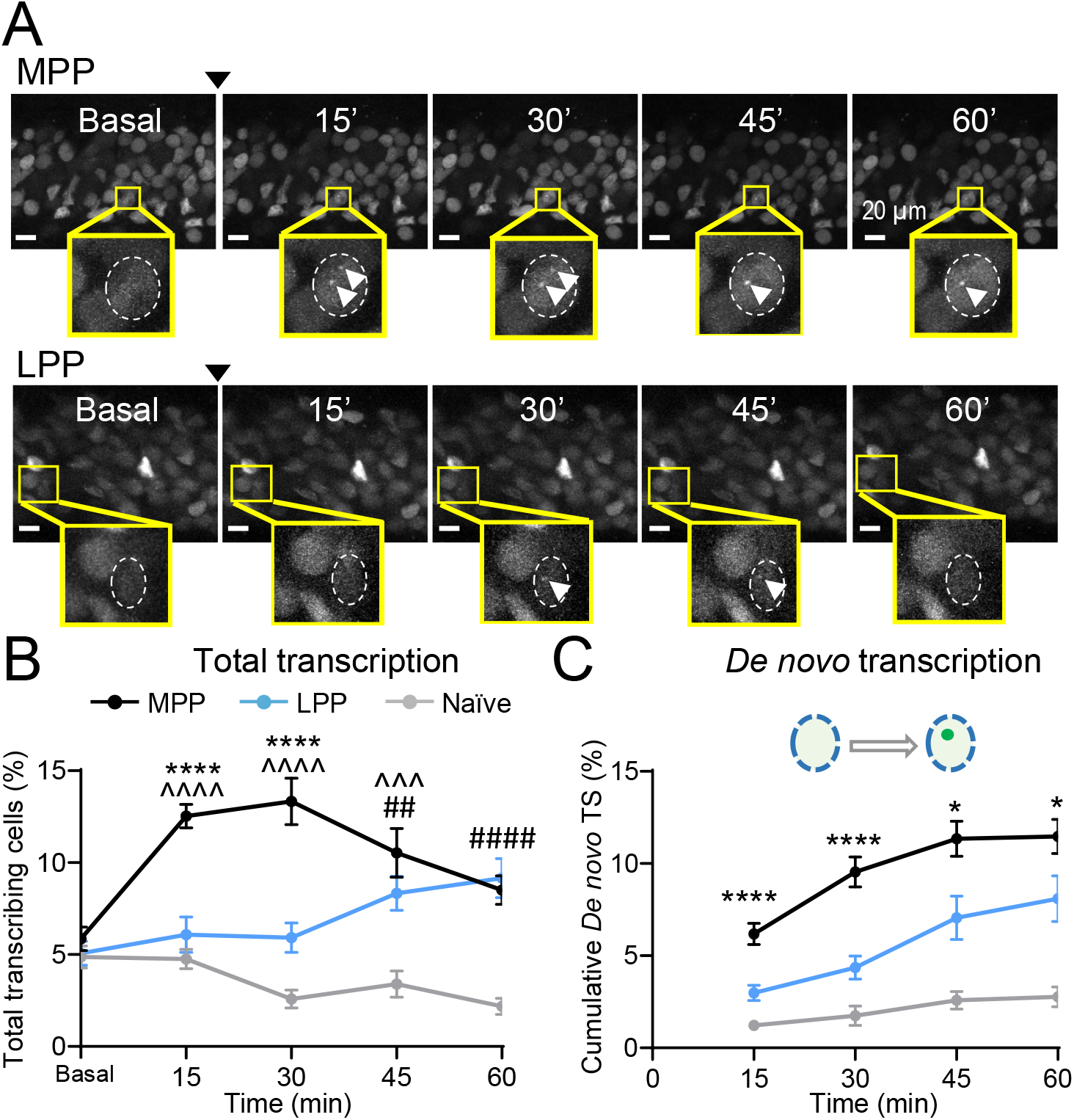
Electrical stimulation of MPP inputs activates *Arc* transcription with rapid onset as compared to LPP activation. (***A***) Representative time course images of MPP *(top)* and LPP *(bottom)* before and after electrical stimulation (denoted by arrowhead). Yellow boxes highlight a GC exhibiting *Arc* transcription in the field of view. (***B***) Quantification of total transcribing cells revealed a significant increase following MPP stimulation (MPP at different time points vs. basal at 0 min; ^^^^ p < 0.001 at 15 min and 30 min; ^^^ p = 0.002 at 45 min; one-way ANOVA). LPP stimulation led to a significant increase in total transcribing cells at later time points (LPP at different time points vs. basal at 0 min; ^##^ p = 0.007 at 45 min, ^####^ p < 0.001 at 60 min; one-way ANOVA). Early time points of transcriptional activation are significantly different between MPP and LPP (MPP vs. LPP at 15 min and 30 min, **** p < 0.001, unpaired *t*-test). Naïve slices showed no increase in transcription signals. (***C***) MPP elicited larger cumulative *de novo* transcription as compared to LPP (MPP vs. LPP at 15 min and 30 min, **** p < 0.001; at 45 min and 60 min, * p < 0.05, unpaired *t*-test). In panels B-C, naïve non-stimulated slices served as control. MPP: n = 9 slices, 7 animals; LPP: n = 8 slices, 6 animals; Naïve: n = 6 slices, 5 animals. Data are presented as mean ± s.e.m. ****, ^^^^, ^####^ denotes p < 0.001; ^^^ denotes p < 0.005, ^##^ denotes p < 0.01, * denotes p < 0.05. Scale bar in images is 20 μm.

To gain deeper insight into activity-dependent induction of *Arc* gene, we selectively quantified GCs displaying *de novo* transcription. While MPP stimulation elicited immediate early *de novo* transcription, a significant delay was observed for LPP (Fig. 2C: MPP 9.5 ± 0.8 % vs. LPP 4.3 ± 0.6 % at 30 min). *De novo* transcription was seen in LPP at 45 min; however, it remained significantly lower than MPP (Fig. 2C). Together, these observations strongly suggested that LPP inputs trigger *Arc* transcription with lower efficacy and longer latency than MPP inputs. Weaker recruitment of LPP inputs or a slower synapse-to-nucleus signal could explain differences in temporal kinetics. To address this possibility, we increased the number of stimulation bursts that may enhance synapse-to-nucleus signaling (34). However, augmenting LPP stimulation in this manner, which increased charge transfer by 3-fold, did not accelerate *Arc* transcription (Fig. S2). The proximity of MPP inputs to GC somata could elicit action potentials and thereby influence *Arc* induction kinetics. We excluded this possibility by performing whole-cell current-clamp recordings that revealed repetitive MPP synaptic activity did not trigger action potentials (Fig. S3). Altogether, our observations suggest that the temporal dynamics of *Arc* transcription display input-specificity.

### Action potential activity of GCs elicits *Arc* transcription

The role of action potential activity in excitation-transcription coupling remains poorly understood (21). Recent evidence indicated that specific forms of neuronal depolarizations impact transcription and translation of selective target genes and proteins, many of which are immediate early in nature (19, 22). Using optogenetic stimulation of GCs, we elicited action potentials and visualized the temporal kinetics of *Arc* transcription (Fig. 3A). Quantification of transcription signals after optogenetic stimulation showed that *Arc* was induced within 15 min, with peak values 12.4 ± 1.6 % at 30 min, and lasted up to 60 min (Fig. 3B). To ensure that ChIEF expression did not result in aberrant upregulation of *Arc* transcription, we delivered antidromic electrical stimulation to GCs and observed comparable immediate early kinetics (Fig. 3B,C). However, a significant difference in transcriptional decay was noted at 45 min (Fig. 3B). Additionally, we detected changes in amplitude of transcription signals with optogenetic and synaptic stimulation (Fig. S4). Based on intensity traces of *de novo* sites, we calculated that activity-induced transcription was maintained for an average of 30 min (Fig. S4), corroborating the ON-duration measured in cultured hippocampal neurons (28). Such observation suggests that the kinetics of the transcriptionally active state may be conserved and intrinsically regulated. Our results established optogenetic stimulation as an effective method to investigate *Arc* transcription in tissue.

**Fig 3.**
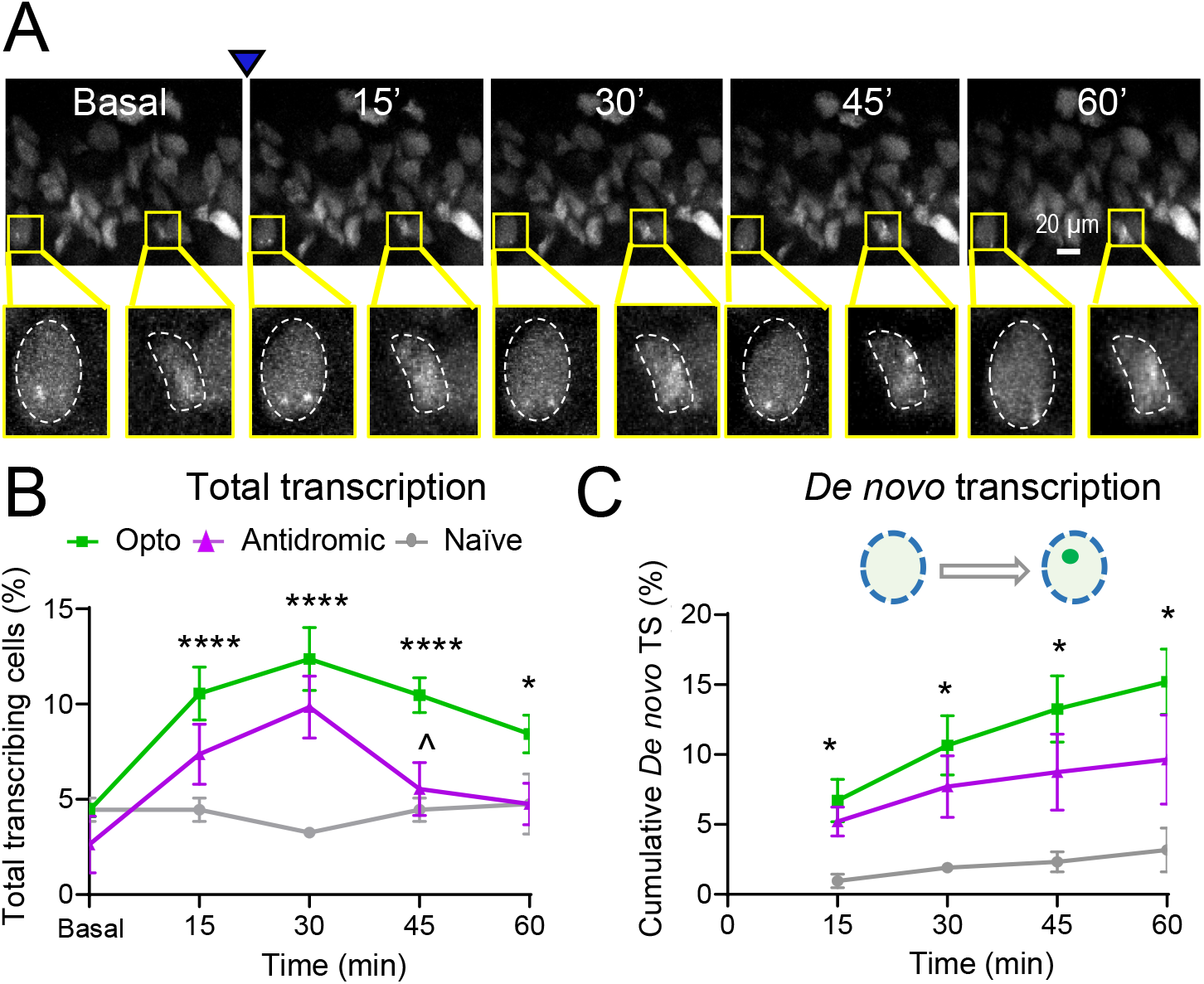
Optogenetic stimulation of GCs induces *Arc* transcription. (***A***) Representative time course images before and after optogenetic stimulation with 473nm light (denoted by arrowhead). Yellow box highlights a GC exhibiting *Arc* transcription in the field of view. (***B***) Quantification of total transcribing cells after optogenetic stimulation showed significant *Arc* activation as compared to baseline (Opto vs. basal at 0 min, **** p < 0.001 at 15 min, 30 min, 45 min; * p = 0.02 at 60 min, one-way ANOVA). Antidromic electrical stimulation of GCs elicits similar *Arc* transcriptional onset as optogenetic activation, however a significant difference in decay is noted at 45 min (Opto vs. Antidromic, ^ p = 0.02 at 45 min, unpaired *t*-test). (***C***) Quantification of *de novo* transcription following optogenetic stimulation revealed a significant increase as compared to naïve slices (Opto vs. Naïve all time points, p = 0.01, unpaired *t*-test). Optogenetic and antidromic stimulations display comparable *de novo* transcription. Opto: n = 8 slices, 6 animals. Naïve: n = 3 slices, 3 animals. Antidromic: n = 3 slices, 3 animals. Data are presented as mean ± s.e.m. * denotes p < 0.05, ** denotes p < 0.01, *** denotes p < 0.005.

### Role of ionotropic glutamate receptors and L-type VGCC in *Arc* transcription

Neuronal activity can trigger Ca^2+^ influx via different postsynaptic ionotropic glutamate receptors (iGluRs), such as AMPA and NMDA receptors, or voltage gated Ca^2+^ channels (VGCCs). Therefore, we determined the impact of these Ca^2+^ sources on the timing and efficacy of *Arc* transcription. MPP stimulation in the presence of the AMPAR antagonist, NBQX (10 μM) and NMDAR antagonist, D-APV (25 μM) significantly reduced *Arc* transcription as compared to control (Fig. 4A, top: MPP 13.3 ± 1.3 %; D-APV+NBQX 3.8 ± 1.1 % at 30 min). The L-type VGCC has been implicated in IEG expression mechanisms (35–37), and is presumably localized to the soma and proximal dendrites of GCs (38). L-type VGCC blockade with nimodipine (30 μM) had no significant effect on immediate early *Arc* transcription induced by MPP stimulation (Fig. 4A, top: MPP 13.3 ± 1.2 %; Nimodipine 10.6 ± 1.2 at 30 min). However, the L-type VGCC may play a role in sustaining transcriptional activation for longer durations (≥ 45 min) after synaptic stimulation (Fig. 4A, top: MPP 10.5 ± 1.3 %, Nimodipine 6.1 ± 0.9 % at 45 min). *De novo* transcription following MPP stimulation was significantly impaired by inhibiting iGluRs (Fig. 4A), but not with L-type VGCC antagonism (Fig. 4A, bottom). Conversely, optogenetic stimulation activated early *Arc* transcription in the presence of iGluR antagonists, while L-type VGCC blockade caused robust reductions in total and *de novo Arc* transcription (Fig. 4B: Opto 12.4 ± 1.6 %; Nimodipine 6.7 ± 1.5 % at 30 min; Opto *de novo* 10.7 ± 2.1 %; Nimodipine *de novo* 4.3 ± 1.0 % at 30 min). Therefore, the L-type VGCC highly influences *Arc* transcription mediated by somatic activation. These findings suggest that the Ca^2+^ influx via postsynaptic iGluRs and the L-type VGCC has a distinct contribution to the immediate early activation of *Arc* and subsequent transcriptional efficacy.

**Fig. 4.**
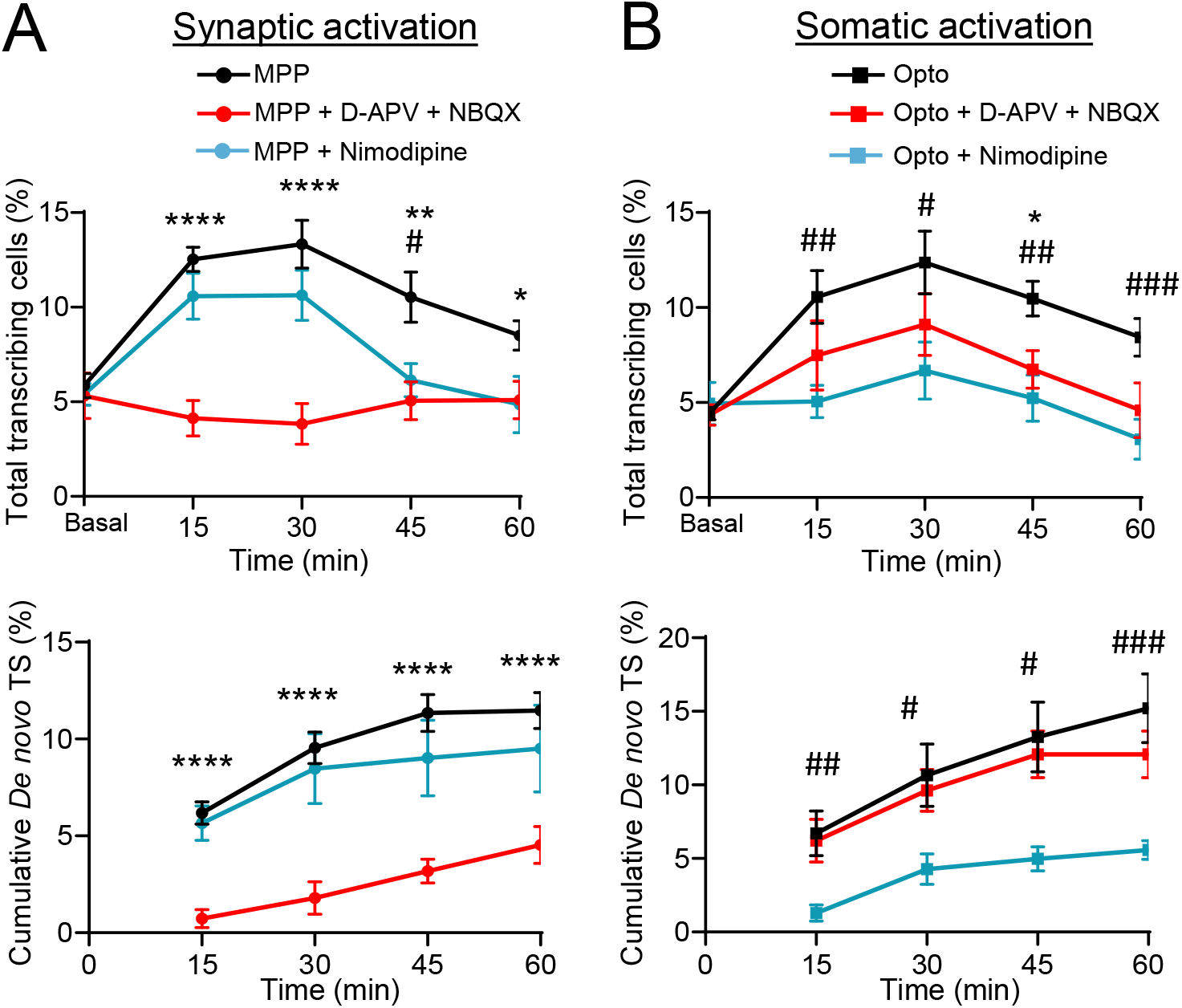
Contribution of iGluRs and L-type VGCC to *Arc* transcription following synaptic or somatic activation. (***A***) Pharmacological inhibition of iGluRs significantly reduces transcriptional activation as compared to control MPP stimulation (MPP vs. NBQX+D-APV, **** p < 0.001 at 15 min, **** p = 0.001 at 30 min, ** p = 0.007 at 45 min, * p = 0.02 at 60 min, unpaired *t*-test). L-type VGCC blockade does not impact early *Arc* induction but impaired maintenance at 45 min (MPP vs. Nimodipine, # p = 0.02 at 45 min, unpaired *t*-test). *De novo* transcription exhibits high sensitivity to iGluR antagonism (MPP vs. NBQX+D-APV, **** p < 0.001, unpaired *t*-test). MPP: n = 9 slices, 7 animals; NBQX + D-APV: n = 5 slices, 4 animals; Nimodipine: n = 6 slices, 5 animals (***B***) L-type VGCC blockade attenuates *Arc* transcription after somatic activation (Opto vs. Nimodipine, ^##^ p = 0.007 at 15 min, ^#^ p = 0.029 at 30 min, ^##^ p = 0.008 at 45 min, ^###^ p = 0.004 at 60 min, unpaired *t*-test). iGluR inhibition significantly impacts *Arc* induction following somatic activation at 45 min (Opto vs. NBQX+D-APV at 45 min, * p = 0.025, unpaired *t*-test). *De novo* transcription is diminished in the presence of Nimodipine following somatic activation (Opto vs. Nimodipine, ^##^ p = 0.009 at 15 min, ^#^ p = 0.02 at 30 min, ^#^ p = 0.01 at 45 min, ^###^ p = 0.004 at 60 min, unpaired *t*-test). Opto: n = 8 slices, 6 animals; NBQX + D-APV: n = 4 slices, 4 animals; Nimodipine: n = 5 slices, 4 animals. Data are presented as mean ± s.e.m. **** denotes p < 0.001, ^###^ denotes p < 0.005, ^##^ ** denotes p < 0.01, ^#^*denotes p < 0.05.

### The endoplasmic reticulum facilitates *Arc* transcription following MPP stimulation

The subcellular mechanisms supporting synapse-to-nucleus communication can deeply influence adaptations in gene expression (35, 39–41). To test the involvement of ER-mediated Ca^2+^ wave propagation (42, 43), we depleted intracellular Ca^2+^ stores using the inhibitor of Ca^2+^-dependent ATPases cyclopiazonic acid (CPA, 30 μM) (43). CPA strongly attenuated both total and *de novo Arc* transcription after MPP stimulation (Fig. 5A: MPP 12.5 ± 0.6 %; CPA 6.8 ± 1.1 % at 15 min; MPP *de novo* 6.2 ± 0.6 %; CPA *de novo* 1.5 ± 0.5 % at 15 min). Moreover, CPA delayed the onset of *de novo* transcription that remained significantly lower than control (Fig. 5A, bottom). In contrast, CPA had no significant effect on total and *de novo Arc* transcription induced by optogenetic activation (Fig. 5B: Opto 10.6 ± 1.4 %; CPA 9.3 ± 2.1 %; Opto *de novo* 6.7± 1.5 %; CPA *de novo* 5.7 ± 2.0% at 15 min). Together, these findings strongly suggest that the ER supports a synapse-to-nucleus signal to promote rapid *Arc* transcription during repetitive MPP activity. In contrast, direct activation of GCs that elicits action potentials induces *Arc* transcription that is independent of the ER.

**Fig. 5.**
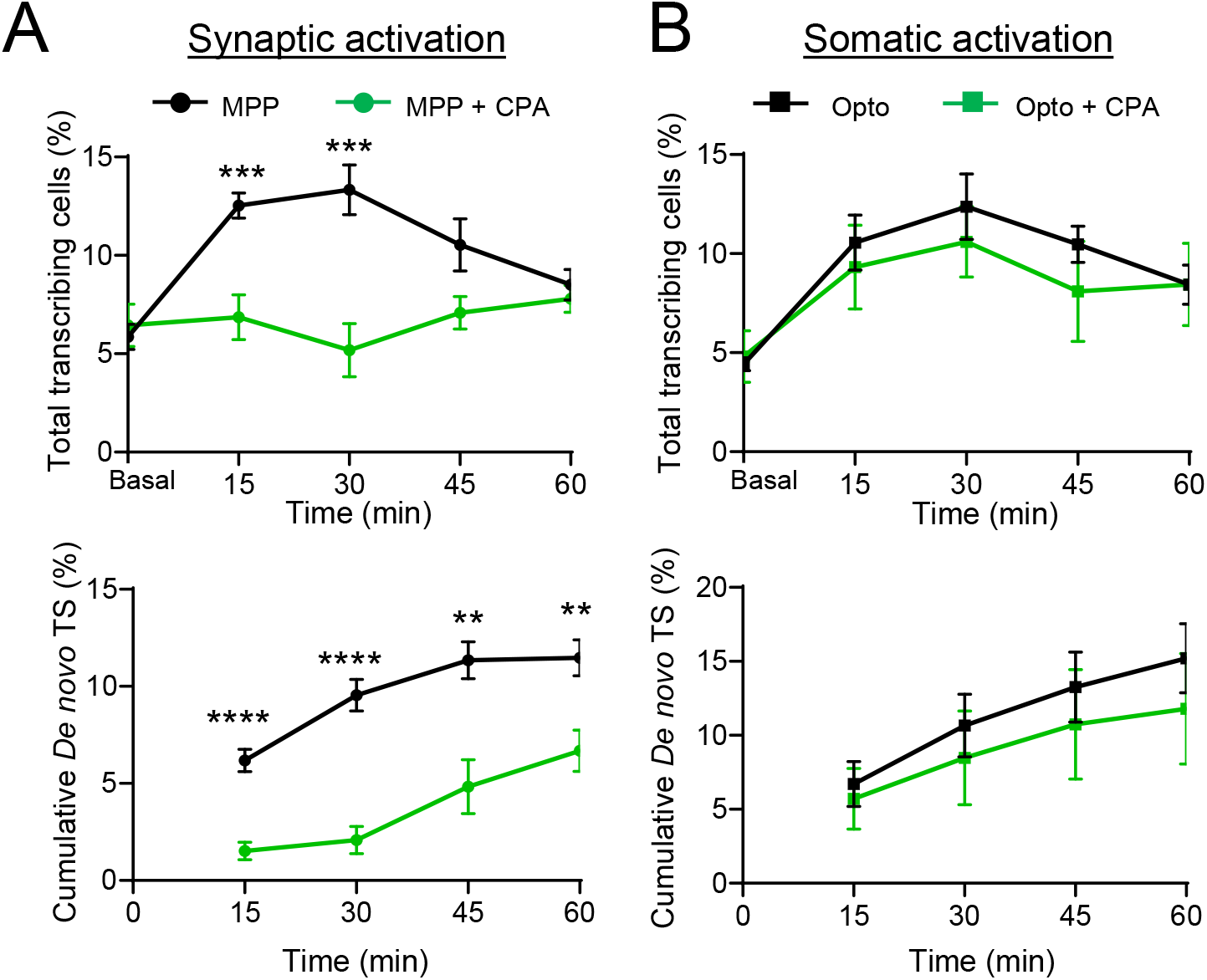
The endoplasmic reticulum supports immediate early *Arc* transcription after MPP stimulation. (***A***) Bath-application of cyclopiazonic acid (CPA, 30 μM) blocks early phase transcription following MPP stimulation (MPP vs. CPA, *** p = 0.003 at 15 min, *** p = 0.001 at 30 min, unpaired *t*-test). CPA treatment attenuates *de novo* transcription (MPP vs. CPA, **** p < 0.001 at 15 min and 30 min, ** p = 0.005 at 45 min, ** p = 0.007 at 60 min, unpaired *t*-test). MPP: n = 9 slices, 7 animals; CPA: n = 5 slices, 5 animals. (***B***) Total and *de novo* transcription are not impacted by CPA treatment after somatic activation. Opto: n = 8 slices, 6 animals; CPA: n = 4 slices, 4 animals. Data are presented as mean ± s.e.m. **** denotes p < 0.001, *** denotes p < 0.005, ** denotes p < 0.01, * denotes p < 0.05.

### MPP inputs elicit nuclear Ca^2+^ elevations supported by the ER

Our observations raised the possibility of ER mediated-Ca^2+^ wave propagation as a synapse-to-nucleus signal supporting *Arc* induction (44–46). To detect nuclear Ca^2+^ elevations in GCs, we expressed a genetically encoded Ca^2+^ indicator containing a nuclear localization signal, NLS-GCaMP6s (Fig. 6A,B). Two weeks post-surgery, acute slices were prepared and stimulation of MPP inputs elicited nuclear Ca^2+^ rise (Fig. 6C). Subsequent bath-application of CPA (30 uM) diminished nuclear Ca^2+^ levels following MPP activity (Fig. 6C: MPP 5.6 ± 1.1, CPA 2.9 ± 0.6; total nuclear Ca^2+^). To discard the possibility that Ca^2+^ reductions were due to photobleaching or repetitive MPP stimulation, we performed successive MPP activations and observed higher Ca^2+^ rise after the second stimulation (Fig. 6D: MPP-1^st^ 4.3 ± 0.9; MPP-2^nd^ 6.6 ± 1.6; total nuclear Ca^2+^). Hence, nuclear Ca^2+^ elevation is supported by the ER and possibly underlies rapid transcription initiation following MPP activation. Given LPP stimulation led to slower transcription kinetics we sought to determine whether this observation can be explained by the nuclear Ca^2+^ levels induced by LPP. Recruitment of LPP inputs resulted in low nuclear Ca^2+^ levels while a subsequent MPP activation significantly increased total Ca^2+^ in the same nuclei (Fig. 6E: LPP 2.1 ± 0.5; MPP 4.7 ± 1.3; total nuclear Ca^2+^). Together our findings indicate GC excitatory synapses distinctly elevate nuclear Ca^2+^ levels. Specifically, MPP inputs elicit large Ca^2+^ elevations in GC nuclei that are supported by the ER.

**Fig. 6.**
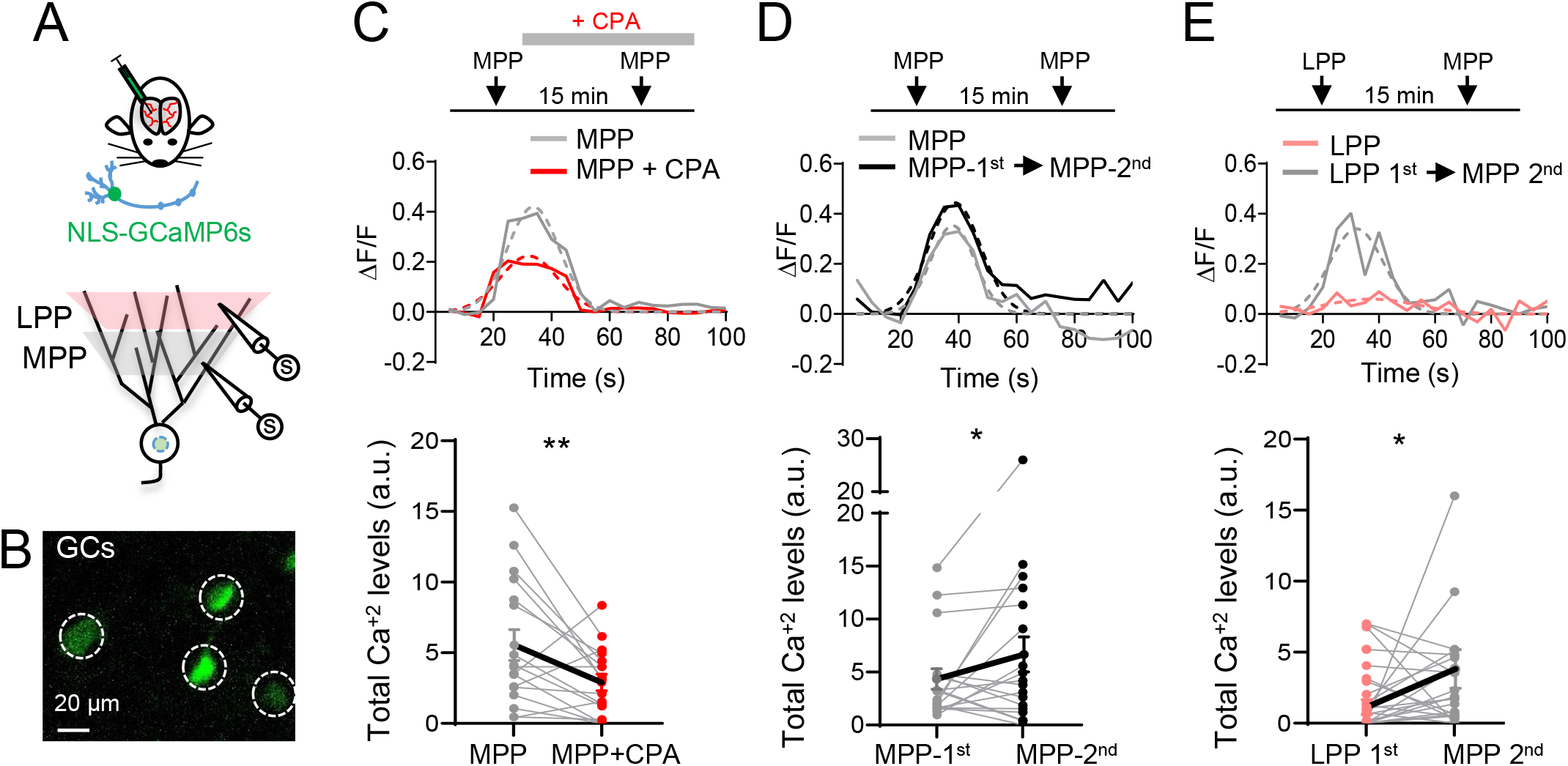
The endoplasmic reticulum supports nuclear Ca^2+^ elevations largely driven by MPP synaptic stimulation. (***A***) The genetically encoded Ca^2+^ indicator containing a nuclear localization signal (NLS-GCaMP6s) was injected into the dentate gyrus of Arc^P/P^/PCP-GFP mice. Two-weeks post-surgery acute hippocampal slices were prepared for the electrical stimulation of perforant path inputs. (***B***) Image depicting the expression of the NLS-GCaMP6s in granule cell (GC) somata. In panels C-E, the experimental design schematic is represented at the top, line profiles of calcium signals are displayed in the middle, and quantification of total nuclear Ca^2+^ at the bottom (each circle indicates individual nuclei, bold line shows the average). (***C***) CPA treatment (30 μM) significantly reduced nuclear Ca^2+^ rise from baseline MPP stimulation (MPP vs. CPA, **p = 0.007, paired *t*-test, 17 ROIs; n = 6 slices, 4 animals). (***D***) Interleaved experiments revealed nuclear Ca^2+^ increases when MPP inputs are activated repetitively (MPP-1^st^ vs. MPP-2^nd^, * p = 0.04, paired *t*-test, 18 ROIs; n = 7 slices, 5 animals). (***E***) Stimulation of MPP inputs after LPP activation results in significantly larger nuclear Ca^2+^ signals (LPP vs. MPP, * p = 0.03, paired *t*-test, 24 ROIs; n = 6 slices, 4 animals,). Bold lines indicate mean ± s.e.m. * denotes p < 0.05, ** denotes p < 0.01.

## Discussion

In this study, we performed high-resolution imaging of *Arc* transcription in real time to elucidate how activity at different spatial locations induces distinct excitation-transcription coupling to regulate *Arc* gene expression in live tissue. We have characterized the impact of different Ca^2+^ sources, iGluRs, L-type VGCC, and the ER, on *Arc* transcription following activation of GCs. Moreover, we have established how specific synaptic inputs (MPP versus LPP) elicit *Arc* transcription with different efficacy and latency. The input-specific temporal kinetics potentially reflects distinct synapse-to-nucleus signaling mechanisms underlying *Arc* activation. In this regard, we uncovered an MPP synapse-to-nucleus signal mediated by the ER that supports large nuclear Ca^2+^ elevations in GCs. Our results highlight the complex relationship of neuronal activity, nuclear Ca^2+^ and gene regulation. Importantly, the temporal differences in subcellular processes that gate IEG transcription in response to GC activity may critically impact hippocampal circuit functions such as learning and memory.

Long-term potentiation, epileptic forms of activity, and memory-related behaviors have been shown to elicit *Arc* transcription in the dentate gyrus of the rodent hippocampus (7–9, 47, 48). Specifically, *in situ* detection of mRNAs has linked *Arc* transcription to animal behaviors including spatial exploration and gustatory memory (23, 49). However, the ability to persistently track transcription of IEGs in the same neuron in tissue had remained a technical challenge in the field. The development of transgenic mice expressing GFP and dVenus under the *Arc* promoter (24–26) enabled *in vivo* monitoring of promoter activity (50), but provided an indirect readout of transcription. By tagging the endogenous *Arc* gene, we showed activity-induced transcription with single allele resolution and detect modes of transcription (amplitude, duration, and *de novo* activation) in live tissue. Employing electrical and optogenetic stimulation of GCs, we establish the ability to delineate how neuronal activity along the somatodendritic axis influences immediate early *Arc* transcription dynamics. Furthermore, the optogenetic approach may be applied in future work to attain insights into action potential driven IEG regulation *in vivo*. The impact of modifying the temporal kinetics of transcription on synaptic plasticity and brain circuit functions remains enigmatic. Future longitudinal studies using this Arc^P/P^-PCP-GFP mouse may provide a deeper understanding of IEG dysregulation in maladaptive behaviors associated with brain disorders such as drug addiction, epilepsy, and cognitive decline.

Neuronal activity occurring throughout the somatodendritic axis can differentially impact IEG expression in neurons (21, 35, 40). Recent work on the IEG *Npas4* proposed that neuronal depolarizations in the form of action potentials and excitatory postsynaptic potentials triggered distinct genomic regulation by NPAS4-heterodimers in CA1 pyramidal neurons (19). Although IEGs display commonalities, including activation without the requirement of protein synthesis, each IEG exhibits precise temporal windows of expression (22, 51). In case of *Arc* transcription in the dentate gyrus, our findings revealed similar temporal dynamics following synaptic or somatic activation, although the underlying signaling mechanisms differed (Figs. 4 and 5). Strikingly, synaptic stimulation along the dendritic axis displayed input-specific *Arc* expression. MPP activation resulted in robust and rapid *Arc* transcriptional onset as compared to LPP (Fig. 2). Differences in the latency and magnitude of *Arc* transcription indicate that structural and functional synaptic properties may determine how the same presynaptic activity transduces distinct transcriptional output. Although our results do not fully discard other possibilities, such as differences in synaptic distance from the soma, the fact that increasing LPP burst stimulation did not alter the onset of *Arc* suggests that synaptic properties likely influence the threshold for IEG activation. Considering that MPP and LPP mainly convey information about context and content of animal experience, respectively (30–33), our findings may implicate how environmental information processed at different synaptic inputs is imprinted in the genome to support hippocampal circuit functions.

Synaptic signals are transmitted to the nucleus through several subcellular processes and employ different sources of Ca^2+^ (21, 35, 40, 41, 44). Our results indicate that iGluRs significantly contribute to *Arc* gene activation (Figure 4), possibly by recruiting signaling cascades and/or by engaging the ER to elevate Ca^2+^ levels in the nucleus to facilitate *Arc* transcription. Inhibition of ER Ca^2+^ mobilization during MPP stimulation exhibited strong attenuation in *Arc* transcription and a significant delay in onset time (Fig. 5). These findings showcase the role of the ER in excitation-transcription coupling after synaptic but not somatic activation (Fig. 5). A similar mechanism of Ca^2+^-induced Ca^2+^ release from the internal stores via ryanodine receptors has recently been reported to upregulate IEGs like *Npas4* (52). In addition, the delayed *de novo* transcription of *Arc* in the presence of CPA, suggests that an alternative slower synapse-to-nucleus mechanism may exist, including activity-dependent shuttling of transcription factors or other proteins from the synapse to the nucleus (41, 53).

MPP stimulation robustly activated *Arc* transcription and generated large nuclear Ca^2+^ elevations, as compared to LPP. Therefore, lower Ca^2+^ rise in the nucleus likely results in slower *Arc* transcriptional activation, as seen following LPP stimulation. Given the synaptic distance of LPP inputs from the GC soma, distal depolarizations may attenuate due to cable properties (54). Recently, NMDAR mediated plateau potentials and sodium dendritic spikes were shown to robustly impact synaptic plasticity of LPP inputs (55). While dendritic Ca^2+^ spike propagation to the soma can activate the transcription factor *NFAT* (56), whether LPP dendritic spike mechanisms compensate for electrical attenuation to support *Arc* transcription remains to be explored. Nonetheless, our results suggest that MPP and LPP could initiate distinct subcellular processes resulting in different temporal regulation of IEGs to support synaptic plasticity of specific GC inputs. Dissecting the molecular and cellular signaling intricacies of LPP and MPP inputs will further our understanding of how environmental features of content and context is conveyed to the dentate gyrus (30–33).

Our study highlights the importance of revealing the underpinnings of IEG activation that may critically impact brain function. In particular, the possibility of studying IEGs in persistent neuronal populations over time provides opportunities to delve into how compartment and input-specific activity regulates gene expression in the mammalian brain. While the induction threshold for IEGs like *Arc*, *Npas4*, *c-fos*, and *zif-268* may vary as previously suggested (20), our findings indicate IEG temporal dynamics may be influenced not only by activity but also in an input specific manner. The precise neuronal activity required to set in motion specific gene programs in different cell types warrants further investigation. Given the implication of IEG dysfunction in brain disorders (45, 57), understanding the temporal kinetics and requirements of IEG expression and their effect on neuronal physiology may provide insights to identify novel therapeutic targets.

## Materials and Methods

### Arc^P/P^ x PCP-GFP transgenic mouse generation

Targeting construct for PCP-GFP was produced by using conventional cloning approaches with homology arms amplified from the C57BL/6J mouse bacterial artificial chromosome vector RPCI-23-81J22. A neocassette containing CAG-Pr-stop/flox-NLS-PCP-GFP was generated and targeted in the ROSA 26 locus. After verification by sequencing, the targeting construct was electroporated into hybrid ES cell line from (ROSA)26, ES cell clones were screened for homologous recombination by PCR. Correctly targeted ES cells were microinjected into blastocysts and chimeras were mated to FLPeR mice (ROSA26::FLP) to remove the FRT-flanked neomycin resistance cassette in the offspring. Pups without the PGK-neocassette were identified by PCR and used to backcross to a C57BL/6J background. These mice contain NLS-PCP-GFP gene inserted into the (ROSA)26 locus, termed PCP-GFP mice. Expression of the PCP-GFP gene is blocked by a DIO loxP-flanked STOP fragment placed upstream NLS-PCP-GFP sequence and driven by the CAG promoter. Successful excision by Cre recombinase is indicated by the nuclear expression of PCP-GFP expression in Cre-expressing tissues. The PCP-GFP mouse was then crossed with the Arc^P/P^ mouse (28) to generate the Arc^P/P^ x PCP-GFP double homozygous mouse. Animal care and experimental procedures were carried out in accordance with the protocols approved by the Institutional Animal Care and Use Committee at Albert Einstein College of Medicine, and Janelia Research Campus of Howard Hughes Medical Institute (HHMI).

### Genotyping

To genotype PCP-GFP R26 lines, PCRs were performed for the following primer sets. R26 wt forward primer (5′-CCAAAGTCGCTCTGAGTTGT-3′), and reverse primers, R26 wt (5′-CCAGGTTAGCCTTTAAGCCT-3′), and CMV R1 (5′-CGGGCCATTTACCGTAAGTT-3′), yielding a 250bp product for the WT allele, and a 329 bp product for the PCP-GFP respectively. Arc-PBS mice were genotyped as described before (Das et al, 2018). Genotyping was performed for both the PBS-KI and R26-PCP-GFP to identify the Arc^P/P^ x PCP-GFP double homozygous mice. PCR conditions were 94°C for 30 s, 55°C for 30 s, and 72°C for 30 s for 35 cycles.

### Stereotaxic Surgery

Arc^P/P^ x PCP-GFP mice at postnatal day 27 (P27) were anesthetized with oxygen-isoflurane flowing at 1.5 ml/min and positioned into a Kopf stereotax instrument. A beveled Hamilton syringe injected 1-1.5 μl of AAV5-CaMKII-mcherry-Cre virus at a rate of 0.15 μl/min at coordinates targeting the dentate gyrus (−2.1 mm A/P, 1.7 mm M/L, 2.5 mm D/V). Two-weeks post-surgery mice were sacrificed for combined electrophysiology and two-photon microscopy. For optogenetic experiments a 1:2 mix of AAV5-CaMKII-mcherry-Cre/AAV-DJ-FLEX-ChIEF-tdTomato virus was injected into the dentate gyrus and animals were sacrificed 5-6 weeks post-surgery for experimentation.

### Hippocampal slice preparation

Arc^P/P^ x PCP-GFP mice > P41 were perfused with 20 mL of cold NMDG solution containing in (mM): 93 NMDG, 2.5 KCl, 1.25 NaH_2_PO_4_, 30 NaHCO_3_, 20 HEPES, 25 glucose, 5 sodium ascorbate, 2 Thiourea, 3 sodium pyruvate, 10 MgCl_2_, 0.5 CaCl_2_, brought to pH 7.35 with HCl. The hippocampi were isolated and cut (300 μm) using a VT1200s microslicer in cold NMDG solution. Acute hippocampal slices were placed in a chamber containing artificial cerebral spinal fluid solution (ACSF) incubated in a warm water bath 33-34°C. The ACSF solution was composed of (in mM): 124 NaCl, 2.5 KCl, 26 NaHCO_3_, 1 NaH_2_PO_4_, 2.5 CaCl_2_, 1.3 MgSO_4_ and 10 glucose. All solutions were equilibrated with 95% O_2_ and 5% CO_2_ (pH 7.4). Post-sectioning, acute slices were allowed to recover at room temperature for at least 45 min prior to experiments. All animals were anesthetized using 4% Isoflurane and sacrificed by decapitation without differentiation of sex.

### Electrophysiology

Extracellular field excitatory postsynaptic potentials (fEPSPs) were recorded at 32 ± 1 °C using a patch-type pipette filled with 1 M NaCl in the presence of the GABAa receptor antagonist, picrotoxin (100 μM). A stimulating glass electrode was filled with ACSF and placed in the medial perforant path (MPP) or lateral perforant path (LPP) to activate inputs with an Isoflex stimulating unit (A.M.P.I, 100 μs pulse width duration). A MultiClamp 700B amplifier (Axon Instruments) registered field responses that were filtered at 10 kHz and digitized at 5 kHz. Stimulation and acquisition were controlled with custom software (Igor Pro 6). After identifying MPP or LPP field responses and obtaining a baseline Z-stack of GC images, a modified high frequency stimulation (HFS) protocol consisting of 100 pulses at 200 Hz, repeated 5 times every 5 s was elicited. To detect action potential generation following optical stimulation ChIEF^+^ GCs were patched using a patch-type pipette (3-4 mΩ) containing a potassium methanosulfate internal consisting of in mM: 135 KMeSO_4_, 5 KCl, 1 CaCl_2_, 5 NaOH, 10 HEPES, 5 MgATP, 0.4 Na_3_GTP, 5 EGTA and 10 D-glucose, pH 7.2 (280-285 mOsm). ChIEF^+^ GCs were held at resting membrane potentials ranging from −80 to −70 mV for whole-cell current-clamp recordings.

### Optogenetics

Acute slices showing optimal tdTomato reporter expression (at least 75% of DG fluorescence) were selected for experimentation. The Invitro Ultima 2P microscope (Bruker Corp.) contains a Coherent 473 nm laser path that delivered optical stimulation of 25 pulses at 25 Hz repeated 20 times every 5 s (8 mW, 2-4 ms pulse duration). The stimulation area was specifically defined using customized Mark Point software (Bruker Corp.) and was empirically determined based on *Arc* allele detection in the field of view described below.

### Two-photon microscopy for *Arc* transcription signal detection

An Invitro Ultima 2P microscope (Bruker Corp.) with and Insight Deep See laser tuned to 910-930 nm visualized *Arc* alleles with 512 x 512 pixel resolution using 4 mW laser power measured at the 60X objective (Nikon, 1.0 NA). GCs expressing the PCP-GFP coat protein were imaged at 1X magnification to detect an *Arc* allele signal that was chosen as a region of interest (ROI) for 2X magnification. A Z-stack of 25 μm thickness with 0.5 μm steps was taken to assess baseline *Arc* transcription signals before electrical or optical stimulation. After stimulation, Z-stack images of 25 μm thickness with 0.5 μm steps were acquired every 15 min for 1 hr.

### *Arc* transcription image processing and analysis

Transcription sites were analyzed in a semi-automated manner using particle tracking on Image J, and using the FISH-Quant software on the MATLAB platform (58). Briefly, the images were filtered, and then semi-automated detection of transcription sites was performed based on the maximum intensity pixel and fitted to a 2D-Gaussian using ROIs of 28-32 pixels for each site. The intensity of each transcription site was normalized by the background signal of the nucleus using the same ROI dimension. The normalized intensity value of 1 represents that no transcription sites are detected. An intensity threshold of 10% change (i.e. values 1.1 or higher) was used to designate a transcription site. The same transcription site was followed over time to measure the change in normalized intensity values, and below 1.1 was considered as transcriptional shutdown. The images were background subtracted and subjected to 2D-Gaussian filtering for representation in figures. Quantification of total transcribing cells were performed as follows:

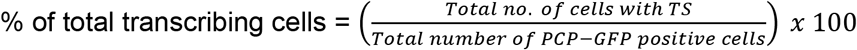

Quantification of *de novo* transcription was performed by determining cells with new transcription sites (1 or 2 alleles) at each time point after stimulation and calculating the cumulative number of cells over time.

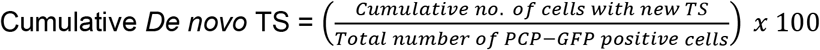

### Two-photon microscopy for nuclear Ca^2+^ imaging

Using the Insight Deep See laser tuned to 945 nm the Invitro Ultimo 2P microscope identified GC somata expressing the GCaMP6s-NLS genetically encoded Ca^2+^ indicator. A stimulating electrode was placed parallel to the identified GCaMP6s^+^ GC somata at MPP or LPP inputs verified by paired pulse stimulation. At 512 x 512 pixel resolution and 1X magnification GC somata were imaged using T-series software to capture calcium elevations in response to electrical burst stimulation (HFS) previously described. Twenty frames were captured at 1 frame/5.01 s containing a baseline period of 4 frames followed by electrical stimulation.

### Nuclear Ca^2+^ imaging analysis

For nuclear Ca^2+^ measurements, images from the time series were Z-projected, and the ROIs for the nucleus were detected in a semi-automated manner on Fiji. The same cells were imaged in first and second stimulation paradigm and the same ROIs with required correction for x-y drift were used to measure nuclear GCaMP fluorescence. The average value from the first 3 frames was considered to be baseline fluorescence (F_0_), and the following change in fluorescence (ΔF) was measured. The values in the traces are represented as ΔF/F. A minimum cutoff of ΔF/F values of 0.05 was used as an inclusion criterion for the ROIs. Traces were fitted to a peak-fitting algorithm and the maximum likelihood analysis was performed for peak assignment. The area under the peak represents the total nuclear Ca^2+^ levels reported.

### Viruses

AAV5-CaMKII-mcherry-Cre virus was obtained from UNC Chapel Hill Vector Core. The DJ-FLEX-ChIEF-tdTomato construct was a generous gift from Dr. Pascal Kaeser and the AAV virus of this construct was custom ordered from UNC Chapel Hill Vector Core. The GCaMP6s-NLS was cloned in house in p323 lentiviral backbone with the GCaMP6s sequences derived from pGP-CMV-GCaMP6s (Addgene plasmid # 40753). High titer lentivirus was generated at the Einstein Virus Core facility.

### Reagents

All chemicals used to prepare cutting and recording solutions were acquired from Millipore Sigma. D-APV, Cyclopiazonic Acid, and Nimodipine, were ordered from Tocris Biosciences. NBQX was obtained from Cayman Chemical Company. Picrotoxin was acquired from Hello Bio Company.

### Data Analysis and Statistics

One-way ANOVA (Dunett’s multiple comparison) was used to determine statistical significance at different time points of transcription imaging compared to baseline (t=0) for total transcription. Paired *t*-test determined statistical significance post-stimulation at different time points for *de novo* transcription quantification across different conditions. Similarly, in Ca^2+^ imaging experiments, paired *t*-test determined statistical significance in before and after pharmacological conditions and repetitive bouts of stimulation. For comparisons in other pharmacological conditions, unpaired *t*-tests were performed. Normality test for all conditions was performed using the Shapiro-Wilk test. Wilcoxon Signed Ranks test was used not normally distributed data. All statistical tests were performed on Graph Pad Prism and Origin Pro 9.

## Acknowledgements

We thank all the Castillo and Singer lab members for invaluable discussions, and Bryen Jordan and Yingxi Lin for critical reading of our manuscript. We thank Caiying Guo and Janelia Gene Targeting and Transgenics Resources for help with the development of the PCP-GFP transgenic mice. We also thank Melissa Lopez Jones for assisting with genotyping and Chiso Nwokafor for animal maintenance. This research was supported by NIH grants R01 NS083085 to R.H.S., R01 MH125772, R01 MH116673, and R01 NS113600 to P.E.C., a Ruth L. Kirschstein NRSA Fellowship F31MH109267 to P.J.L. and R21 MH122961 to S.D.

## Author Contributions

P.J.L. and S.D. designed and analyzed experiments. R.H.S. and P.E.C provided guidance and supervision. P.J.L. performed stereotaxic surgeries, electrophysiology, and two-photon microscopy. SD developed *Arc*^P/P^ x *Arc*^PCP-GFP^ mouse lines and analyzed two-photon imaging data. P.J.L. and S.D. wrote the first manuscript, which was edited by all authors.

---

**Fig. S1.**
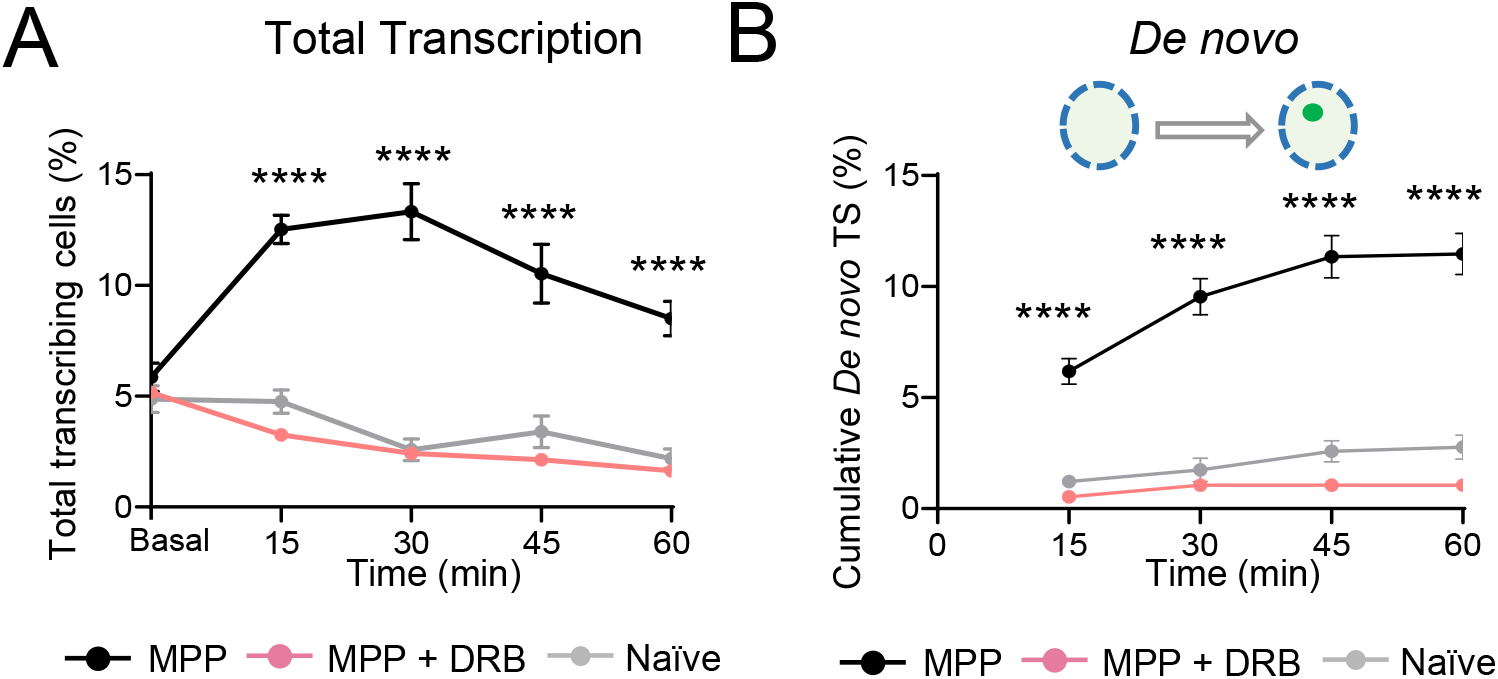
*Arc* activation is prevented using a transcription inhibitor. (***A***) Control slices significant *Arc* upregulation as compared to naïve condition and slices treated with the transcription elongation inhibitor DRB, 100 μM (MPP vs. DRB, **** p < 0.001 at all time points, unpaired *t*-test). *(**B***) *De novo* transcription was not observed in naïve and DRB treated slices as compared to control (MPP vs. DRB, **** p < 0.001 at all time points, unpaired *t*-test). Data are presented as mean ± s.e.m. MPP: n = 9 slices, 7 animals; MPP + DRB: n = 4 slices, 3 animals; Naïve: n = 6 slices, 5 animals. **** denotes p < 0.001

**Fig. S2.**
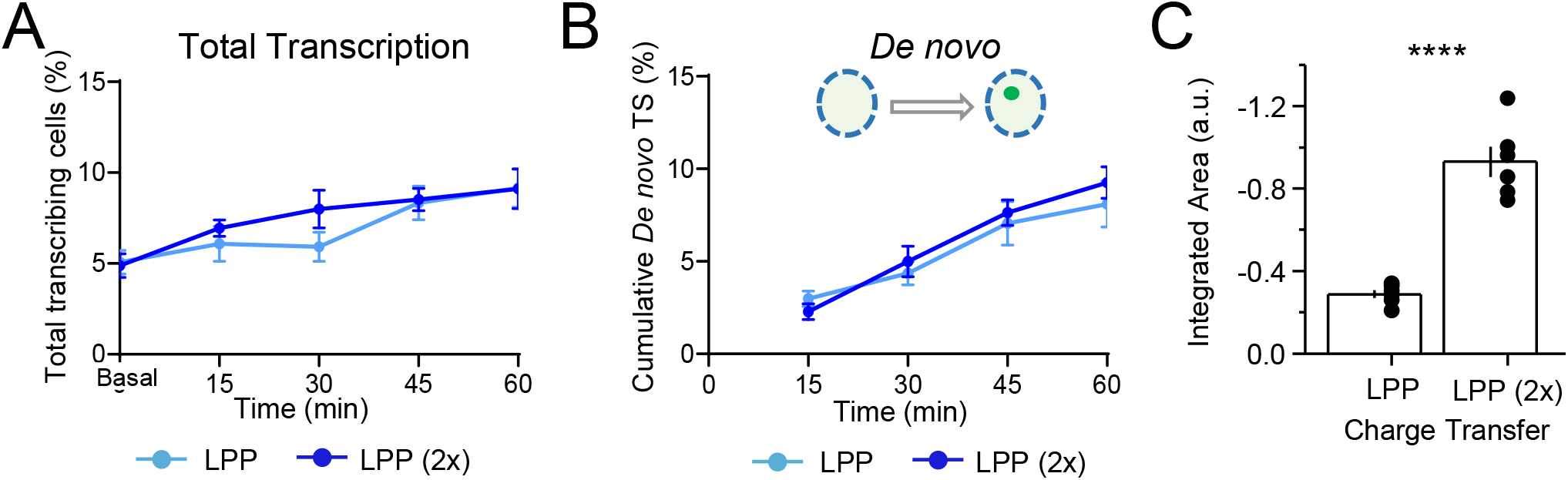
Increasing number of bursts delivered to LPP inputs does not enhance *Arc* activation. ***(A)*** LPP stimulation protocol was repeated 2x in an attempt to enhance *Arc* transcriptional onset. Comparison of total transcriptional of LPP vs. LPP (2x) stimulation showed no significant difference in *Arc* induction (LPP vs. LPP (2x), p > 0.05, unpaired t-test). ***(B)*** Quantification of *de novo* transcription revealed LPP (2x) stimulation activated *Arc* transcription with similar kinetics as LPP protocol. ***(C)*** Integrated area of stimulation protocols revealed larger charge transfer in LPP (2x) condition (LPP: −0.287 ± 0.02, LPP (2x): −0.93 ± 0.07; LPP vs. LPP (2x), p < 0.001, unpaired *t*-test). Data are presented as mean ± s.e.m. LPP: n = 6 slices, 6 animals, LPP (2x): n = 6 slices, 4 animals, **** denotes p < 0.001.

**Fig. S3.**
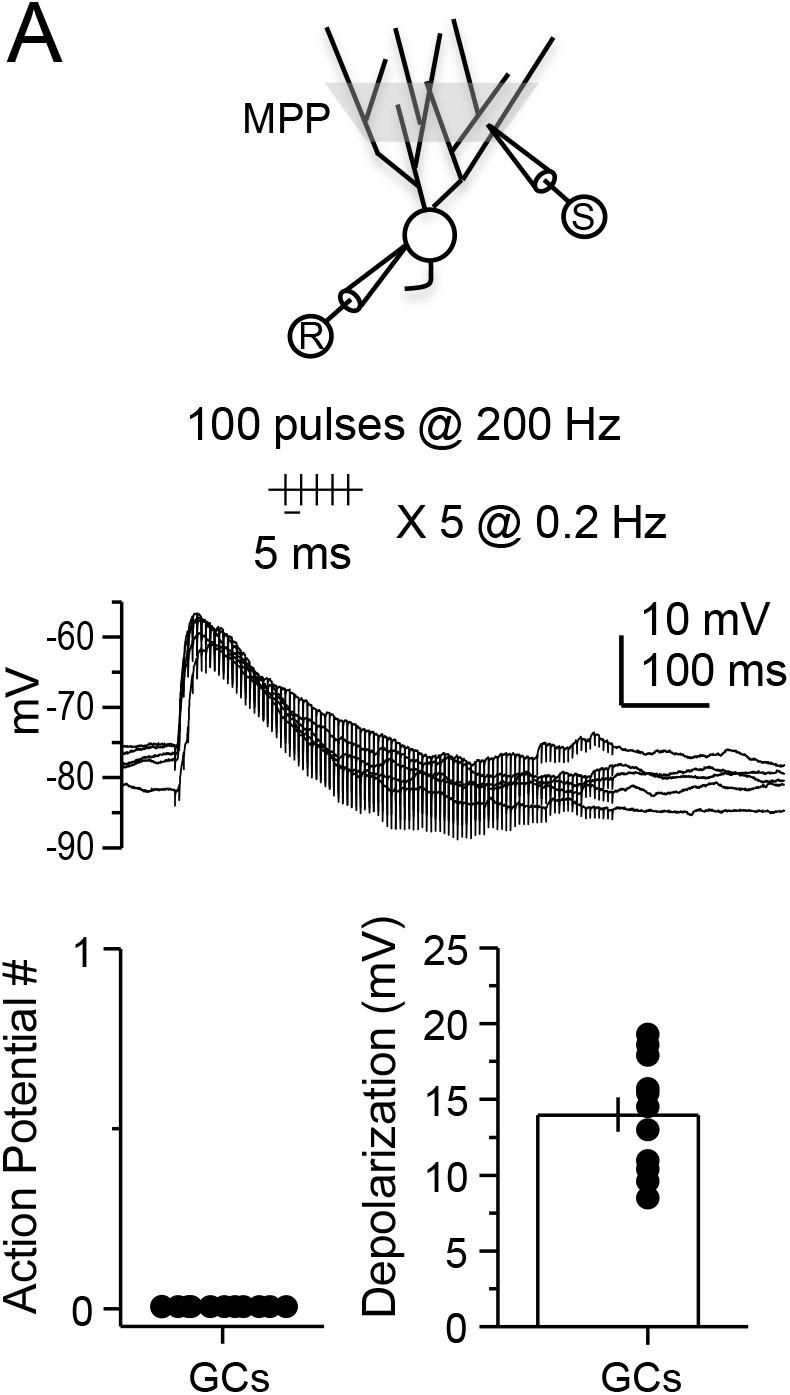
Electrical stimulation protocol of MPP synapses does not elicit action potentials. (***A***) Whole-cell current-clamp recordings of GCs revealed the lack of action potential induction during stimulation protocol of MPP inputs. Representative traces of excitatory postsynaptic potentials during high-frequency stimulations (*middle*). Summary graphs of quantified action potentials (*bottom, left*) and peak depolarization magnitude from stimulation, mV: 13.9 ± 1 (*bottom*, *right*). Data are presented as mean ± s.e.m. n = 11 cells, 6 animals.

**Fig. S4.**
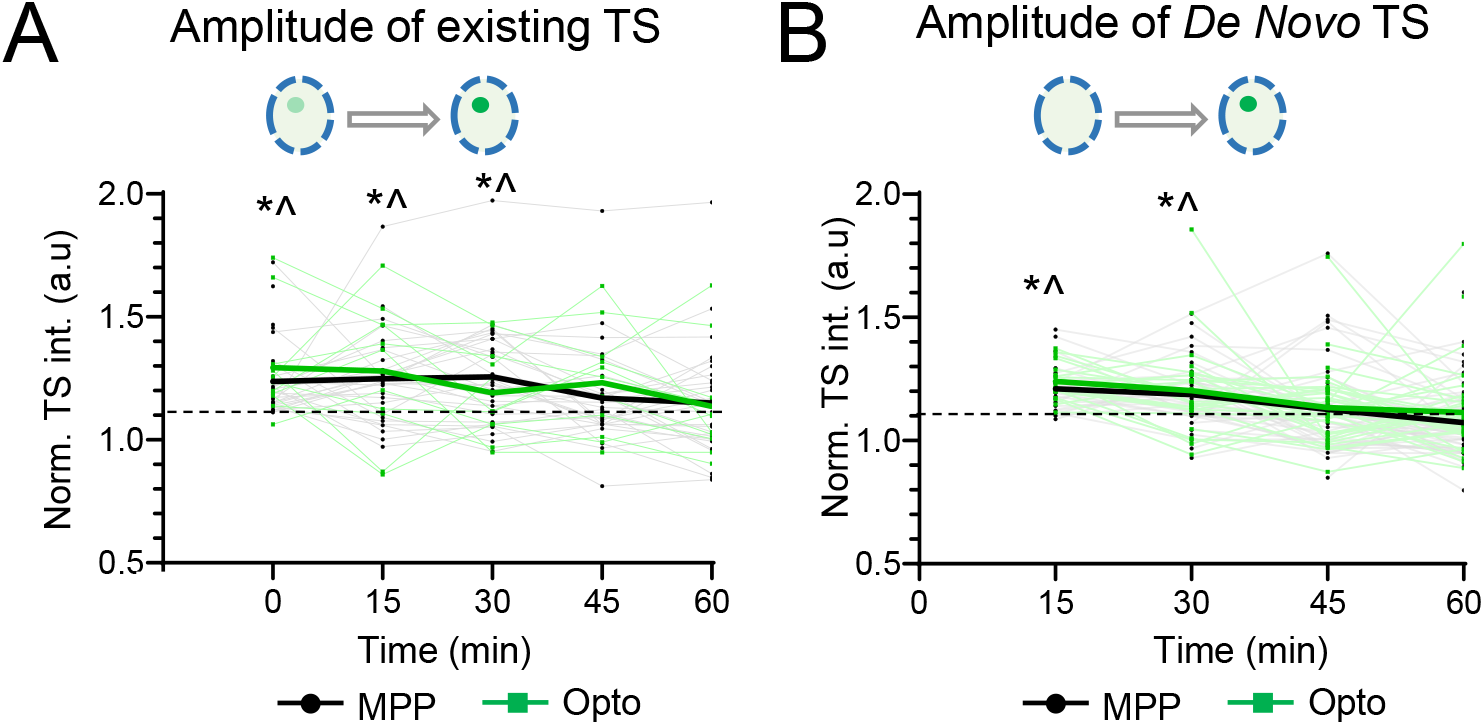
Changes in amplitude of *Arc* transcription following synaptic or somatic activation are detectable. (*A*) Quantification of signal intensity from pre-existing sites following MPP and Opto stimulation reported changes in amplitude of transcriptional output. ***(B)*** *De novo* transcriptional output following MPP and Opto stimulation. In panels A-B, dashed line depicts threshold of > 10% change in amplitude over background to qualify for inclusion. (* p < 0.05 for MPP transcription site above threshold, ^ p < 0.05 for Opto transcription site above threshold, one sample Wilcoxon signed ranks test, n = 40-50 cells per condition). Bold lines indicate the mean response. Data are presented as mean ± s.e.m. * denotes p < 0.05, ^ denotes p < 0.05.

**Movie S1:**
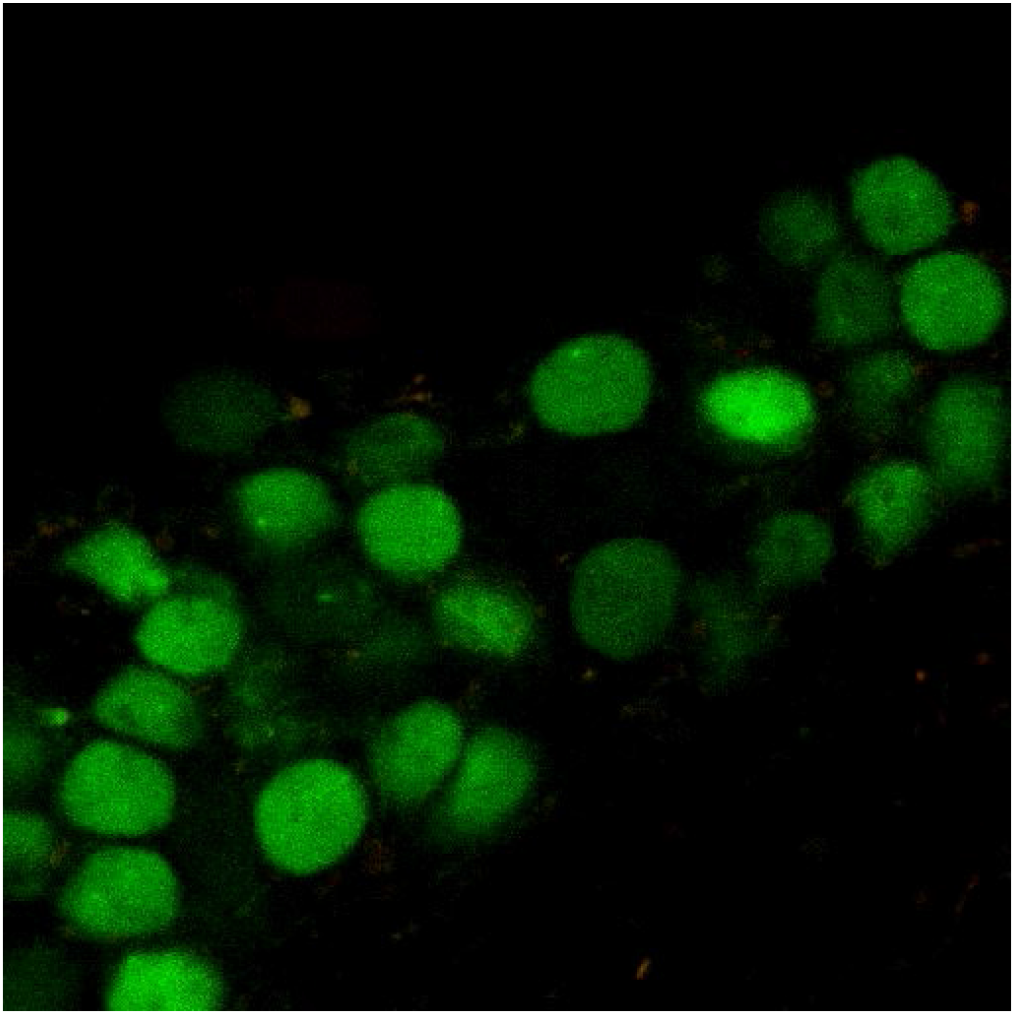
2-photon Imaging *Arc* transcription in acute hippocampal slices. PCP-GFP labeled GC nuclei are seen, where each bright foci indicates a transcribing *Arc* allele. Z-step is 500nm.

## References

1. J. F. Guzowski, B. Setlow, E. K. Wagner, J. L. McGaugh, Experience-dependent gene expression in the rat hippocampus after spatial learning: a comparison of the immediate-early genes Arc, c-fos, and zif268. J Neurosci 21, 5089–5098 (2001).

2. K. Minatohara, M. Akiyoshi, H. Okuno, Role of Immediate-Early Genes in Synaptic Plasticity and Neuronal Ensembles Underlying the Memory Trace. Front Mol Neurosci 8, 78 (2015).

3. V. Ramirez-Amaya et al., Spatial exploration-induced Arc mRNA and protein expression: evidence for selective, network-specific reactivation. J Neurosci 25, 1761–1768 (2005).

4. J. F. Guzowski et al., Inhibition of activity-dependent arc protein expression in the rat hippocampus impairs the maintenance of long-term potentiation and the consolidation of long-term memory. J Neurosci 20, 3993–4001 (2000).

5. N. Plath et al., Arc/Arg3.1 is essential for the consolidation of synaptic plasticity and memories. Neuron 52, 437–444 (2006).

6. J. E. Ploski et al., The activity-regulated cytoskeletal-associated protein (Arc/Arg3.1) is required for memory consolidation of pavlovian fear conditioning in the lateral amygdala. J Neurosci 28, 12383–12395 (2008).

7. O. Steward, C. S. Wallace, G. L. Lyford, P. F. Worley, Synaptic activation causes the mRNA for the IEG Arc to localize selectively near activated postsynaptic sites on dendrites. Neuron 21, 741–751 (1998).

8. W. Link et al., Somatodendritic expression of an immediate early gene is regulated by synaptic activity. Proc Natl Acad Sci U S A 92, 5734–5738 (1995).

9. G. L. Lyford et al., Arc, a growth factor and activity-regulated gene, encodes a novel cytoskeleton-associated protein that is enriched in neuronal dendrites. Neuron 14, 433–445 (1995).

10. S. Chowdhury et al., Arc/Arg3.1 interacts with the endocytic machinery to regulate AMPA receptor trafficking. Neuron 52, 445–459 (2006).

11. E. Messaoudi et al., Sustained Arc/Arg3.1 synthesis controls long-term potentiation consolidation through regulation of local actin polymerization in the dentate gyrus in vivo. J Neurosci 27, 10445–10455 (2007).

12. J. D. Shepherd, M. F. Bear, New views of Arc, a master regulator of synaptic plasticity. Nat Neurosci 14, 279–284 (2011).

13. M. W. Waung, B. E. Pfeiffer, E. D. Nosyreva, J. A. Ronesi, K. M. Huber, Rapid translation of Arc/Arg3.1 selectively mediates mGluR-dependent LTD through persistent increases in AMPAR endocytosis rate. Neuron 59, 84–97 (2008).

14. M. E. Klein et al., Sam68 Enables Metabotropic Glutamate Receptor-Dependent LTD in Distal Dendritic Regions of CA1 Hippocampal Neurons. Cell Rep 29, 1789–1799 e1786 (2019).

15. M. Kyrke-Smith et al., The immediate early gene Arc is not required for hippocampal long-term potentiation. J Neurosci 10.1523/JNEUROSCI.0008-20.2021 (2021).

16. J. D. Shepherd et al., Arc/Arg3.1 mediates homeostatic synaptic scaling of AMPA receptors. Neuron 52, 475–484 (2006).

17. C. L. Wee et al., Nuclear Arc Interacts with the Histone Acetyltransferase Tip60 to Modify H4K12 Acetylation(1,2,3). eNeuro 1(2014).

18. E. Korb, C. L. Wilkinson, R. N. Delgado, K. L. Lovero, S. Finkbeiner, Arc in the nucleus regulates PML-dependent GluA1 transcription and homeostatic plasticity. Nat Neurosci 16, 874–883 (2013).

19. G. S. Brigidi et al., Genomic Decoding of Neuronal Depolarization by Stimulus-Specific NPAS4 Heterodimers. Cell 179, 373–391 e327 (2019).

20. P. F. Worley et al., Thresholds for synaptic activation of transcription factors in hippocampus: correlation with long-term enhancement. J Neurosci 13, 4776–4786 (1993).

21. J. P. Adams, S. M. Dudek, Late-phase long-term potentiation: getting to the nucleus. Nat Rev Neurosci 6, 737–743 (2005).

22. K. M. Tyssowski et al., Different Neuronal Activity Patterns Induce Different Gene Expression Programs. Neuron 98, 530–546 e511 (2018).

23. J. F. Guzowski, B. L. McNaughton, C. A. Barnes, P. F. Worley, Environment-specific expression of the immediate-early gene Arc in hippocampal neuronal ensembles. Nat Neurosci 2, 1120–1124 (1999).

24. M. Eguchi, S. Yamaguchi, In vivo and in vitro visualization of gene expression dynamics over extensive areas of the brain. Neuroimage 44, 1274–1283 (2009).

25. V. Grinevich et al., Fluorescent Arc/Arg3.1 indicator mice: a versatile tool to study brain activity changes in vitro and in vivo. J Neurosci Methods 184, 25–36 (2009).

26. K. H. Wang et al., In vivo two-photon imaging reveals a role of arc in enhancing orientation specificity in visual cortex. Cell 126, 389–402 (2006).

27. H. Sato, S. Das, R. H. Singer, M. Vera, Imaging of DNA and RNA in Living Eukaryotic Cells to Reveal Spatiotemporal Dynamics of Gene Expression. Annu Rev Biochem 89, 159–187 (2020).

28. S. Das, H. C. Moon, R. H. Singer, H. Y. Park, A transgenic mouse for imaging activity-dependent dynamics of endogenous Arc mRNA in live neurons. Sci Adv 4, eaar3448 (2018).

29. J. Y. Lin, M. Z. Lin, P. Steinbach, R. Y. Tsien, Characterization of engineered channelrhodopsin variants with improved properties and kinetics. Biophys J 96, 1803–1814 (2009).

30. H. Lee, D. GoodSmith, J. J. Knierim, Parallel processing streams in the hippocampus. Curr Opin Neurobiol 64, 127–134 (2020).

31. C. Wang, X. Chen, J. J. Knierim, Egocentric and allocentric representations of space in the rodent brain. Curr Opin Neurobiol 60, 12–20 (2020).

32. T. H. Hoang, V. Aliane, D. Manahan-Vaughan, Novel encoding and updating of positional, or directional, spatial cues are processed by distinct hippocampal subfields: Evidence for parallel information processing and the “what” stream. Hippocampus 28, 315–326 (2018).

33. E. S. Nilssen, T. P. Doan, M. J. Nigro, S. Ohara, M. P. Witter, Neurons and networks in the entorhinal cortex: A reappraisal of the lateral and medial entorhinal subdivisions mediating parallel cortical pathways. Hippocampus 29, 1238–1254 (2019).

34. C. P. Bengtson, H. E. Freitag, J. M. Weislogel, H. Bading, Nuclear calcium sensors reveal that repetition of trains of synaptic stimuli boosts nuclear calcium signaling in CA1 pyramidal neurons. Biophys J 99, 4066–4077 (2010).

35. K. Deisseroth, P. G. Mermelstein, H. Xia, R. W. Tsien, Signaling from synapse to nucleus: the logic behind the mechanisms. Curr Opin Neurobiol 13, 354–365 (2003).

36. T. H. Murphy, P. F. Worley, J. M. Baraban, L-type voltage-sensitive calcium channels mediate synaptic activation of immediate early genes. Neuron 7, 625–635 (1991).

37. J. I. Morgan, T. Curran, Role of ion flux in the control of c-fos expression. Nature 322, 552–555 (1986).

38. J. W. Hell et al., Identification and differential subcellular localization of the neuronal class C and class D L-type calcium channel alpha 1 subunits. J Cell Biol 123, 949–962 (1993).

39. T. Kawashima et al., Synaptic activity-responsive element in the Arc/Arg3.1 promoter essential for synapse-to-nucleus signaling in activated neurons. Proc Natl Acad Sci U S A 106, 316–321 (2009).

40. T. H. Ch′ng, K. C. Martin, Synapse-to-nucleus signaling. Curr Opin Neurobiol 21, 345–352 (2011).

41. B. A. Jordan, M. R. Kreutz, Nucleocytoplasmic protein shuttling: the direct route in synapse-to-nucleus signaling. Trends Neurosci 32, 392–401 (2009).

42. S. Watanabe, M. Hong, N. Lasser-Ross, W. N. Ross, Modulation of calcium wave propagation in the dendrites and to the soma of rat hippocampal pyramidal neurons. J Physiol 575, 455–468 (2006).

43. A. Verkhratsky, Physiology and pathophysiology of the calcium store in the endoplasmic reticulum of neurons. Physiol Rev 85, 201–279 (2005).

44. H. Bading, Nuclear calcium signalling in the regulation of brain function. Nat Rev Neurosci 14, 593–608 (2013).

45. E. L. Yap, M. E. Greenberg, Activity-Regulated Transcription: Bridging the Gap between Neural Activity and Behavior. Neuron 100, 330–348 (2018).

46. G. E. Hardingham, F. J. Arnold, H. Bading, Nuclear calcium signaling controls CREB-mediated gene expression triggered by synaptic activity. Nat Neurosci 4, 261–267 (2001).

47. M. K. Chawla et al., Sparse, environmentally selective expression of Arc RNA in the upper blade of the rodent fascia dentata by brief spatial experience. Hippocampus 15, 579–586 (2005).

48. A. Vazdarjanova et al., Spatial exploration induces ARC, a plasticity-related immediate-early gene, only in calcium/calmodulin-dependent protein kinase II-positive principal excitatory and inhibitory neurons of the rat forebrain. J Comp Neurol 498, 317–329 (2006).

49. S. N. Burke et al., Differential encoding of behavior and spatial context in deep and superficial layers of the neocortex. Neuron 45, 667–674 (2005).

50. V. Jakkamsetti et al., Experience-induced Arc/Arg3.1 primes CA1 pyramidal neurons for metabotropic glutamate receptor-dependent long-term synaptic depression. Neuron 80, 72–79 (2013).

51. A. E. West, M. E. Greenberg, Neuronal activity-regulated gene transcription in synapse development and cognitive function. Cold Spring Harb Perspect Biol 3(2011).

52. P. Lobos et al., RyR-mediated Ca(2+) release elicited by neuronal activity induces nuclear Ca(2+) signals, CREB phosphorylation, and Npas4/RyR2 expression. Proc Natl Acad Sci U S A 118(2021).

53. S. Zhai, E. D. Ark, P. Parra-Bueno, R. Yasuda, Long-distance integration of nuclear ERK signaling triggered by activation of a few dendritic spines. Science 342, 1107–1111 (2013).

54. R. Krueppel, S. Remy, H. Beck, Dendritic integration in hippocampal dentate granule cells. Neuron 71, 512–528 (2011).

55. S. Kim, Y. Kim, S. H. Lee, W. K. Ho, Dendritic spikes in hippocampal granule cells are necessary for long-term potentiation at the perforant path synapse. Elife 7(2018).

56. A. R. Wild et al., Synapse-to-Nucleus Communication through NFAT Is Mediated by L-type Ca(2+) Channel Ca(2+) Spike Propagation to the Soma. Cell Rep 26, 3537–3550 e3534 (2019).

57. F. T. Gallo, C. Katche, J. F. Morici, J. H. Medina, N. V. Weisstaub, Immediate Early Genes, Memory and Psychiatric Disorders: Focus on c-Fos, Egr1 and Arc. Front Behav Neurosci 12, 79 (2018).

58. F. Mueller et al., FISH-quant: automatic counting of transcripts in 3D FISH images. Nat Methods 10, 277–278 (2013).

